# Allosteric Logic Gate

**DOI:** 10.64898/2026.04.29.721721

**Authors:** Davide Cois, Sahand Jamal Rahi, Paolo De Los Rios

## Abstract

Allostery enables proteins to transmit local perturbations to distant functional residues, providing a biophysical basis for molecular signal integration. Here we introduce an *Allosteric Logic Gate* (ALG): an elastic network designed to convert two independent deformations at input sites into a Boolean-like conformational output at the distant active region. We model ligand binding as constrained local deformations at two spatially separated sites and read the output through a conformational measure at the active region. We show that it is possible to optimise the network’s spring constants to produce a triggered allosteric response only when both inputs are present, thereby implementing a Boolean AND gate. Moreover, the evolved networks display a strongly non-linear response, matching the switch-like property of logic gates. Statistical analysis of successful networks reveals conserved mechanical motifs, including stiff bonds connecting the input regions and flanking floppy regions that accommodate the output deformation. These results demonstrate that coarse-grained elastic networks can be inverse-designed to perform non-linear logic operations, suggesting a route toward programmable binary allostery, protein-inspired circuits, and biosensors capable of integrating multiple molecular signals.

## INTRODUCTION

The conformational plasticity of proteins is key to their functions, for example to allow enzymes to adapt to different substrates, bind them tightly and then release the reaction products. Allostery, defined as a localised structural perturbation triggering conformational changes at distant protein sites [1, 2], represents a remarkable evolutionary strategy to control plasticity and further diversify cellular functionality [3–5].

Because of their biological relevance, allosteric proteins, and the principle of allostery itself, have been at the forefront of research over the past decades, from the seminal work by Monod, Wyman, and Changeux (MWC) [1], and by Koshland, Nemethy, and Filmer (KNF) [6], which established early theoretical frameworks describing allosteric transitions between distinct protein states, to sophisticated computational approaches capable of explicitly capturing protein dynamics [7–9]. Not surprisingly, the control of allostery has also attracted interest as a potential therapeutic strategy, for example by targeting the allosteric sites with specifically designed drugs [10]. Presently, allosteric proteins also represent one of the new frontiers in protein design [11–16].

Despite its relevance, however, studying allostery is challenging because the large range of involved timescales makes it computationally demanding using detailed, atomistic-scale models. By trading accuracy for effectiveness, elastic network models (ENMs) have emerged as a powerful tool to capture the essential features of the medium-to-large scale dynamics of proteins around their native state. ENMs describe proteins as networks of masses, corresponding to the amino-acids, interconnected by harmonic springs capturing residue-residue interactions. This coarse-grained method can efficiently extract collective modes directly linked to functionally relevant conformational changes and long-range allosteric signal transduction [17–21].

The basic assumption behind ENMs is that dynamical features encompassing the whole protein in a coordinated way should be captured by the lowest frequency/longer “wavelength” normal modes that, being collective and large-scale in nature, depend on averaged, rather than detailed, interactions [20–25].

Because of their simplicity, ENMs naturally connect protein mechanics to material science, and have been widely used to study engineered “allosteric materials” [26–29]. These systems are networks of masses and springs whose parameters are tuned so that a prescribed deformation in one region (input) produces a desired deformation at a distant site (output). In this context, ENMs have revealed that signal propagation across these networks requires a special organisation of the spring stiffnesses, such that input and output regions are connected by “soft” paths that can support low-energy, long-range elastic modes. The same framework was recently used to study multi-state structural conformations of proteins triggered by ligand binding [30] and to design proteins with switchable conformational states [11, 12, 15]. All these works highlight the engineering potential of allostery.

More broadly, ENMs and related coarse-grained models provide minimal physical descriptions of how local perturbations can be transmitted through a protein or artificial elastic network. In single-basin descriptions, allosteric coupling is mediated by collective soft modes; in multi-state descriptions, ligand binding can instead switch the system between distinct stable conformations [30]. Complementary network-based, evolutionary, and machine-learning approaches have been developed to identify residues and pathways that mediate this long-range communication [31–33]. Here we take a complementary inverse-design perspective: rather than reconstructing an existing allosteric pathway, we ask whether an elastic net-work can be programmed so that two independent distal perturbations are integrated into a prescribed Boolean output.

This previous work inspired us to develop here the *Allosteric Logic Gate* (ALG). A logic gate is a function mapping discrete binary inputs to a discrete binary output, according to a truth table. In the framework of allosteric elastic networks, an ALG corresponds to a network designed to change its conformation only in the presence of precise values of at least two independent inputs, implemented as forced deformations localized in two mutually distal sites that are in turn far away from the network regions where the output is detected. In the present work, by combining ENM-based methods with evolutionary algorithms, we construct an elastic network that not only retains the key trait of allostery with long-range conformational change but also performs the discrete non-linear binary operations characteristic of logic gates, as desired.

This innovative application extends the use of ENMs to logic devices and advances our understanding of allosteric signal transduction, potentially making the design of programmable biological machines and novel biosensors a step closer [34–36].

## THE ALLOSTERIC LOGIC GATE

In our ALG design, the network is constructed from a triangular lattice whose nodes are slightly displaced with respect to the perfect lattice, and the new positions are taken as the equilibrium ones for the springs connecting them (see Methods for details). In turn, the stiffness of each spring is determined by the nature of the “amino-acid” (residue) that are assigned to the corresponding pair of nodes, according to some rules. Here we choose them from four different strengths (see Table I and Fig.1).

**TABLE 1.**
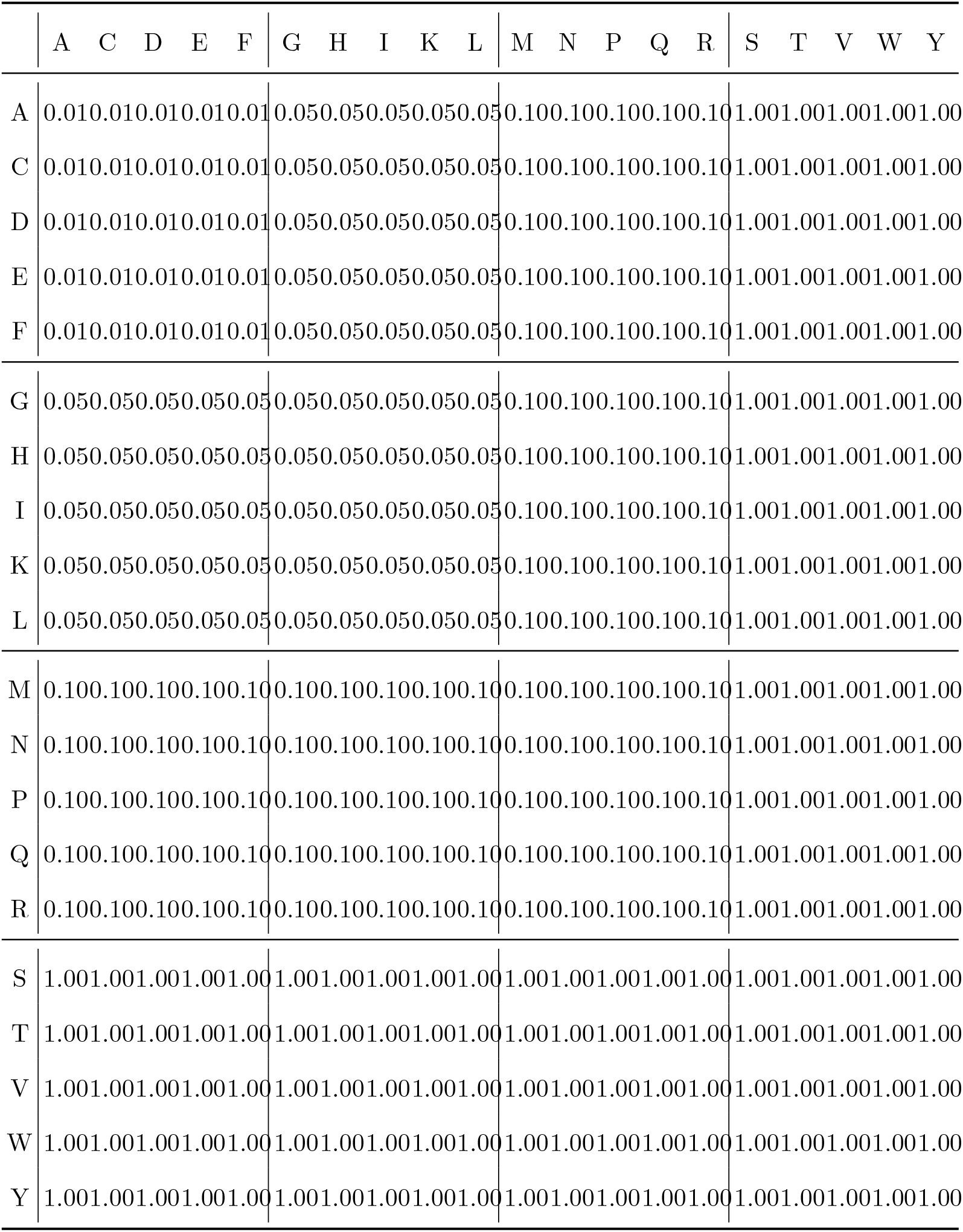
Elastic constant table for amino-acid Interactions. The amino-acid pairings and elastic bond values do not reflect the real chemical bond strengths.

Two pairs of nodes, distant from each other, are designated as *input sites*, or “ligand-binding” sites, and the presence or absence of a ligand is modelled as a forced deformation of those sites (Fig.1). In response to this deformation, the network relaxes to a distorted conformation (not represented in Fig.1), with a consequent structural change in a distant region, the *active site*, whose conformation must change in a required way. Here we read the network response as a sum of distances measured on the first row of the network (Fig. 1), the *allosteric distance, D*_allo_,

**FIG. 1.**
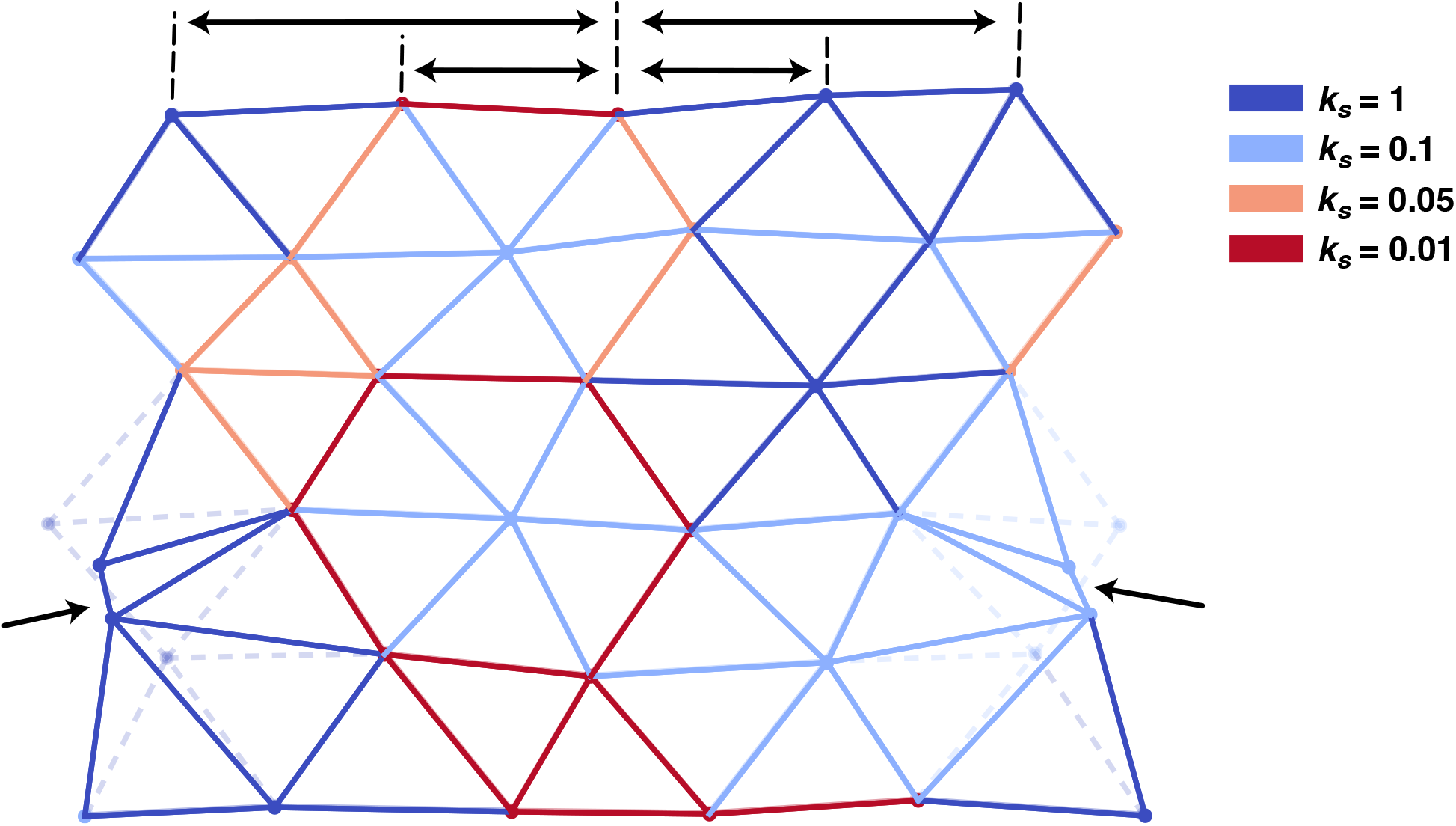
The Elastic Network Model used in this work. A regular triangular lattice is deformed by slightly randomly displacing each node. The springs connecting pairs of nodes can have four different *k*_*s*_. The input deformation is applied on two separated lateral sites (indicated by the arrows; the undeformed network is shown with transparent, dashed lines). The output deformation is related to the distances in the top row, indicated by double arrows.

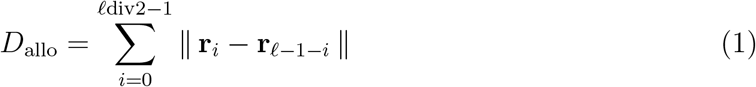

where **r**_*i*_ is the vector of coordinates of the *i*-th node in the active (output) region, *𝓁* is the length of the lattice, *i*.*e*. the number of nodes per row of the network (*e*.*g. 𝓁* = 6 in Fig. 1), *div* is the integer division and ∥ · ∥ denotes the Euclidean norm. In this work, we want *D*_allo_ to be small in response to the correct input.

More specifically, we designed here an ALG that implements a logical AND operation: *D*_allo_ must be small only in the presence of both ligands (*i*.*e*. local perturbations in the binding, input sites), while it should not change (or change as little as possible) if only one, or none, is present. (see Methods for details).

Starting from a random residue distribution, we apply an iterative algorithm to evolve the initial network toward one responding according to the desired AND logic function. To estimate how closely the network implements the AND logics, we used the loss function ℒ

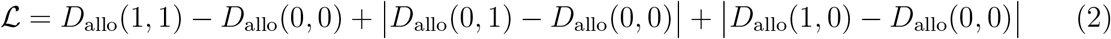

where *D*_allo_(*x, y*) is the allosteric distance in the presence inputs (*x, y*) ∈ {0, 1} × {0, 1} (*x* = 1 when the first allosteric site is bound, *i*.*e*. under constrained displacement, *x* = 0 otherwise, and *y* similarly for the second site. Therefore *D*_allo_(0, 0) is the allosteric distance in the relaxed state of the unperturbed network etc.), and … denotes the absolute value. ℒ is minimised when the allosteric measure of the doubly bound state, (1, 1), is smaller than the one of the unbound state, (0, 0), and the allosteric measures of the singly bound states, (1, 0) and (0, 1), are close to the one of the unbound state, implying that the network does not discriminate between no input and single inputs, as required for an AND gate. We then used the minimization of ℒ as the objective of an iterative, simulated-annealing-like algorithm to guide the network from its random initial conditions toward an optimal ALG configuration reproducing the AND function (see Methods for details).

Due to the intrinsic disorder of the network and to the stochasticity of the algorithm, the solutions that we find, namely the choice of residue in each node, are all different from each other. Some network realisations ranging from allosteric networks with good and strong responses (Figs.2A and B, respectively), to networks almost not responding to the inputs, despite the algorithm having reached the maximum allowed number of iterations(Fig.2C).

**FIG. 2.**
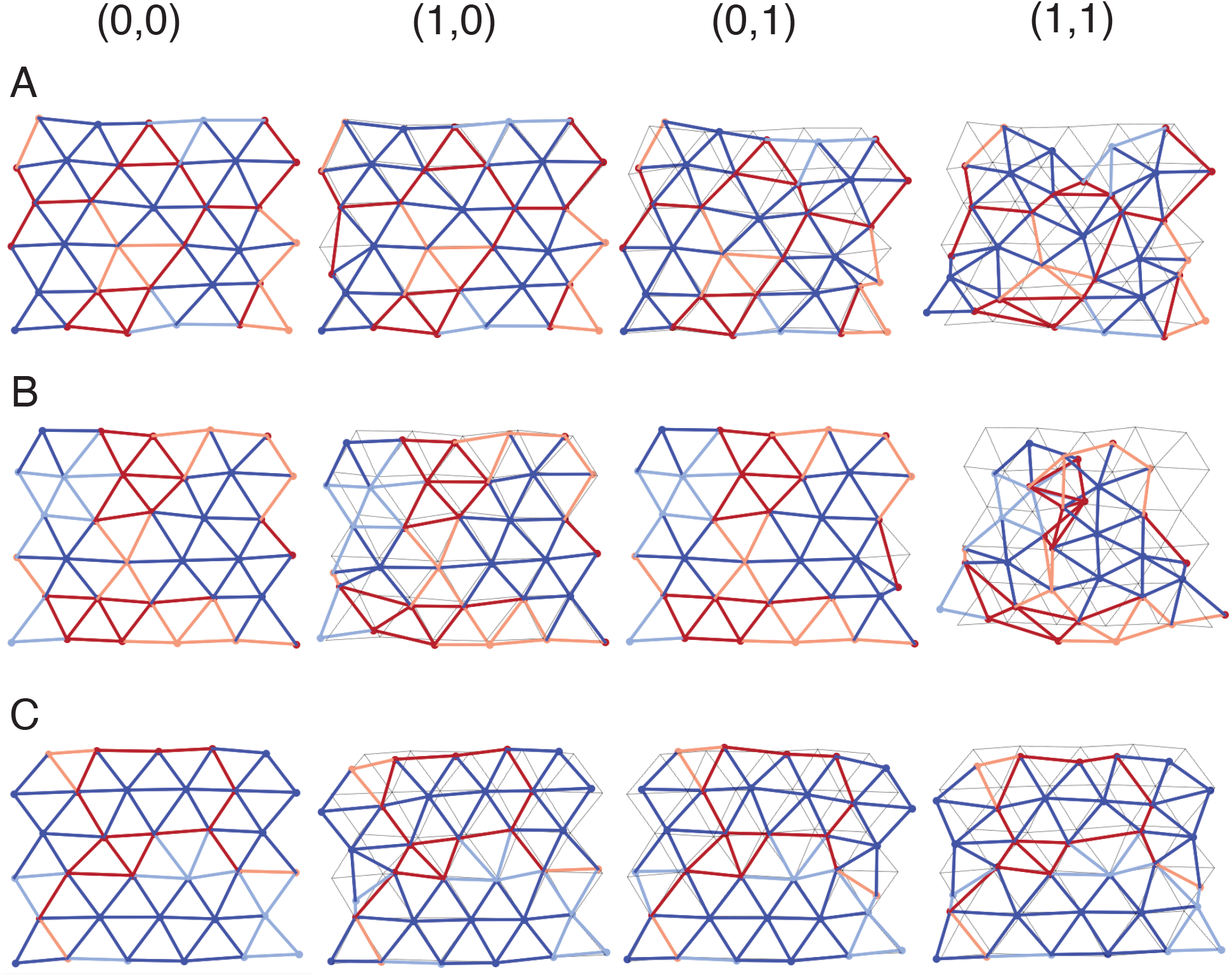
Examples of networks obtained at convergence of the iterative algorithm, with good (A, ℒ= −1.583), strong (B, ℒ = −3.939) and weak (C, ℒ = −0.616) responses. Each row shows the response of the three network for the four different inputs: (0, 0), no deformations; (1, 0), deformation imposed only on the left input site; (0, 1), deformation imposed only on the right input site; (1, 1), deformation imposed on both sites. The bonds are colored according to their stiffness, as from Fig.1 and the undeformed network (no input, first column) is drawn in fainter black lines for reference in the other input cases.

This strong non-linearity, which is indeed required for an AND gate, arises because the springs, which individually follow Hooke’s law with a linear force-extension relation, collectively result in a non-linear response when connected in more than one dimensions.

In this work we analyze networks with *𝓁* = 6. While investigating the behaviour on larger lattices would be desirable, this is impractical for the goal of a thorough exploration of the parameter space, because the convergence time of the algorithm increases roughly quadratically with the system size. Nevertheless, we tested an *𝓁* = 8 network (Figure 3), which likewise showed that an allosteric AND gate can be evolved, albeit with greater computational effort.

**FIG. 3.**
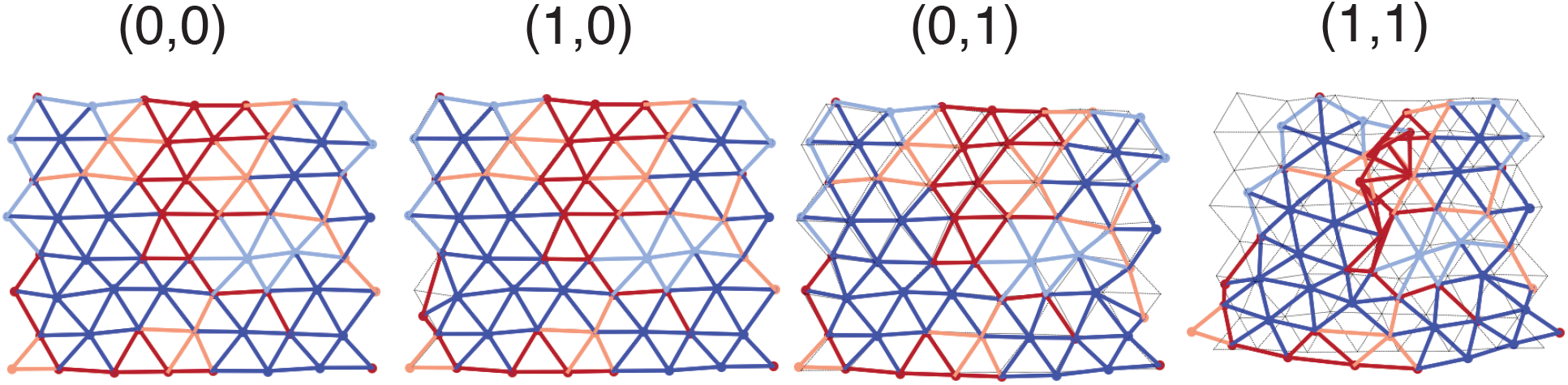
Example of an ALG on a network with 8 nodes per side.

We also investigated the dependence of amplitude of the deformation, as captured by *D*_allo_, on the *magnitude* of the ligand-induced displacement, i.e. by how much the bond of each allosteric site is compressed when the corresponding ligand input is applied. We apply a full deformation to one input and gradually scaled the other input deformation using a parameter *λ* such that *λ* = 0 corresponds to no input and *λ* = 1 to the full input deformation. As seen in Fig.4, the response is highly non-linear in *λ*, remaining close to the one of the single input case (*λ* = 0) up to *λ* ≃ 0.7, then abruptly changing to a new value, that again remains roughly constant up to the full input limit (*λ* = 1). Such non-linearity is precisely what is required by an AND logical function, and more in general by a binary logic gate, and is possible thanks to the non-linear response of a network of linear springs in more than one dimensions.

**FIG. 4.**
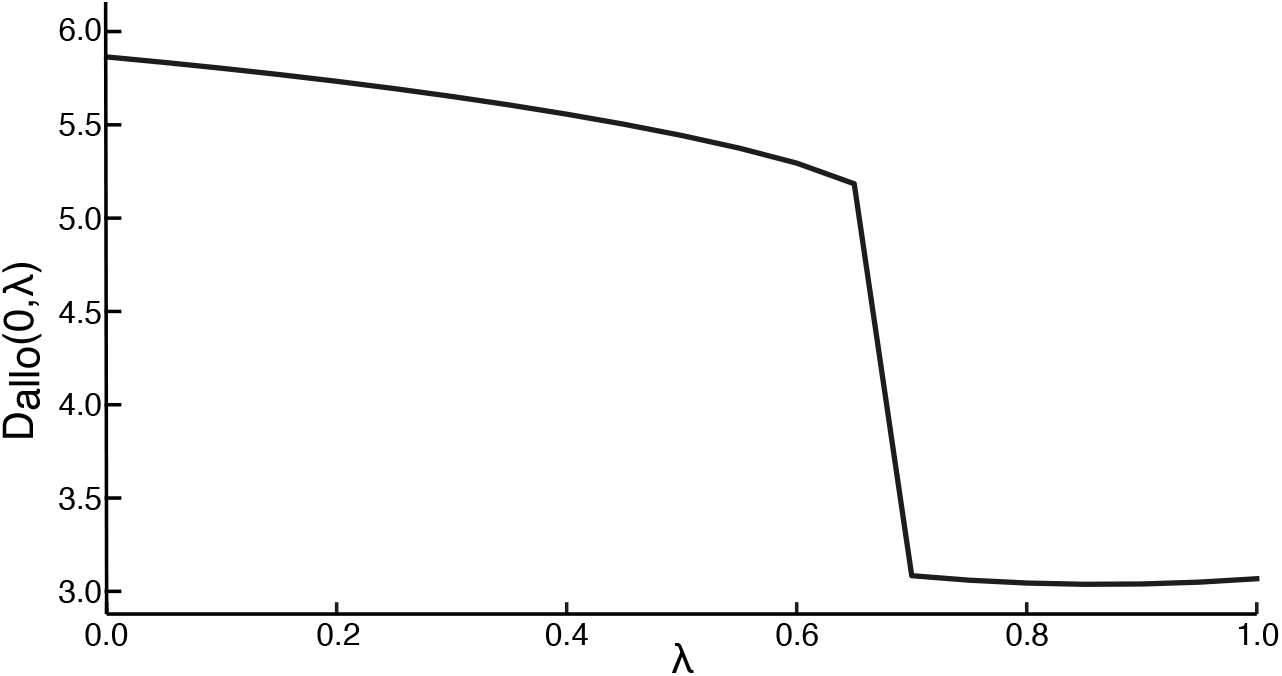
Response function, *D*_allo_(1, *λ*), as a function of *λ*, which controls the amplitude of the deformation imposed on the input site on the right: *λ* = 0, no deformation (*D*_allo_(1, 0)); *λ* = 1, full deformation (*D*_allo_(1, 1)).

## NETWORK ANALYSIS

We selected the evolved networks whose loss function value was below a selected threshold (ℒ *<* −2) to retain the strongest allosteric responses. These networks are characterised by an active region that “crumbles” in response to ligand binding, as shown in Figure 2B.

Out of 66 networks, each with a different lattice noisy initialisation, 19 displayed the crumbling allosteric behaviour, while 47 did not. To investigate the possible reasons behind failures and successes, we *transplanted* the amino-acid sequence from a network that exhibited a significant response onto networks that did not. In no case the sequence transplant produced crumbling in the geometry that did not evolve it on its own (Figure 5A). We then transferred the amino-acid sequences between networks that had *both* crumbling responses. In some cases, this exchange of sequences and nodes coordinates destroyed the crumbling effect (Figure 5B). Thus, there appears to be a tight interplay between the specific node coordinates and the evolved bond configuration that underlies a strong allosteric reaction. Hence, no single “universal” amino-acid configuration guarantees crumbling across all node geometries. This may explain why only about one-third of initialisations reached a strongly negative loss function.

**FIG. 5.**
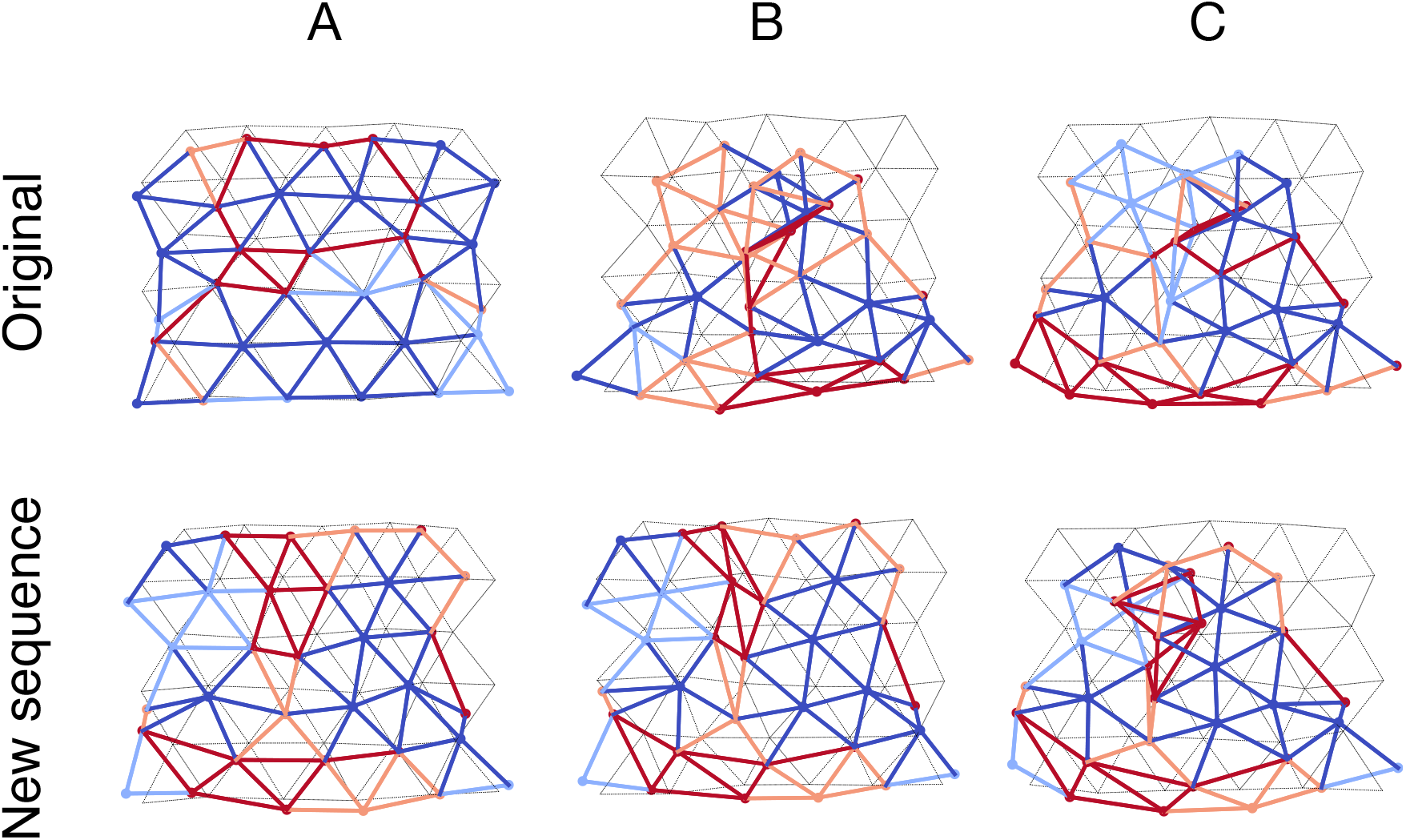
Example of networks with different lattice coordinates, due to different initial noise disordering, with an amino-acid sequence from an allosteric network that evolved a strong (crumbling) response. (A) applied to a network that had an original weak response: no implanted crumbling phenomenon (ℒ_original_ = −0.616 → ℒ_new_ = −0.391). (B) applied to a network that had an original crumbling response: removed crumbling response (ℒ_original_ = −4.415 → ℒ_new_ = −0.834). (C) applied to another network that had an original crumbling response: it keeps the crumbling response (ℒ_original_ = −4.176 → ℒ_new_ = −3.98).

We hypothesise two possible explanations for why the remaining majority did not converge to that solution:

step 1: The evolutionary landscape is complex and rugged. Despite the simulated annealing strategy, most initialisations fail to locate the global minimum of the loss function.

step 2: The emergence of a significant allosteric behaviour strongly depends on the geometry of the network (i.e. the precise node coordinates). Even a relatively small noise (0.1 *𝓁*) of difference between different initialisations of the lattice can significantly alter how the network can evolve a response to ligand binding.

We then performed a statistical analysis of the bond configurations of the ALGs with ℒ *<* −2, investigating how similar the bond configurations were across the different realisations. For each bond, we computed the Shannon entropy of the recorded stiffnesses in the various networks, as defined in Equation 3.

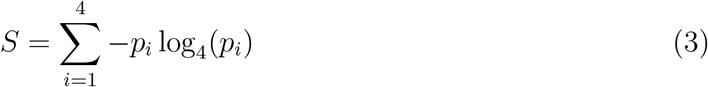

where *p*_*i*_ is the measured frequency of a certain elastic constant of a bond across the ALGs that have been analysed. The entropy measures how consistently for a specific bond an elastic constant is evolved out of the four possible values in order to minimise the loss. *S* = 1 is the function’s maximum and occurs for 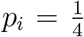, so when all the elastic constants have equal probability, which means that the bond stiffness does not play a significant role in the development of the desired allostery. Conversely, the minimum *S* = 0 occurs for *p*_*i*_ = 1, indicates that the bond always adopts the same stiffness in all the crumbling networks, meaning it plays a crucial role in stabilising or enabling the response. The network of the most conserved bonds in this statistical analysis is shown in Figure 6A, where the bonds are the darkest when their entropy is the lowest.

**FIG. 6.**
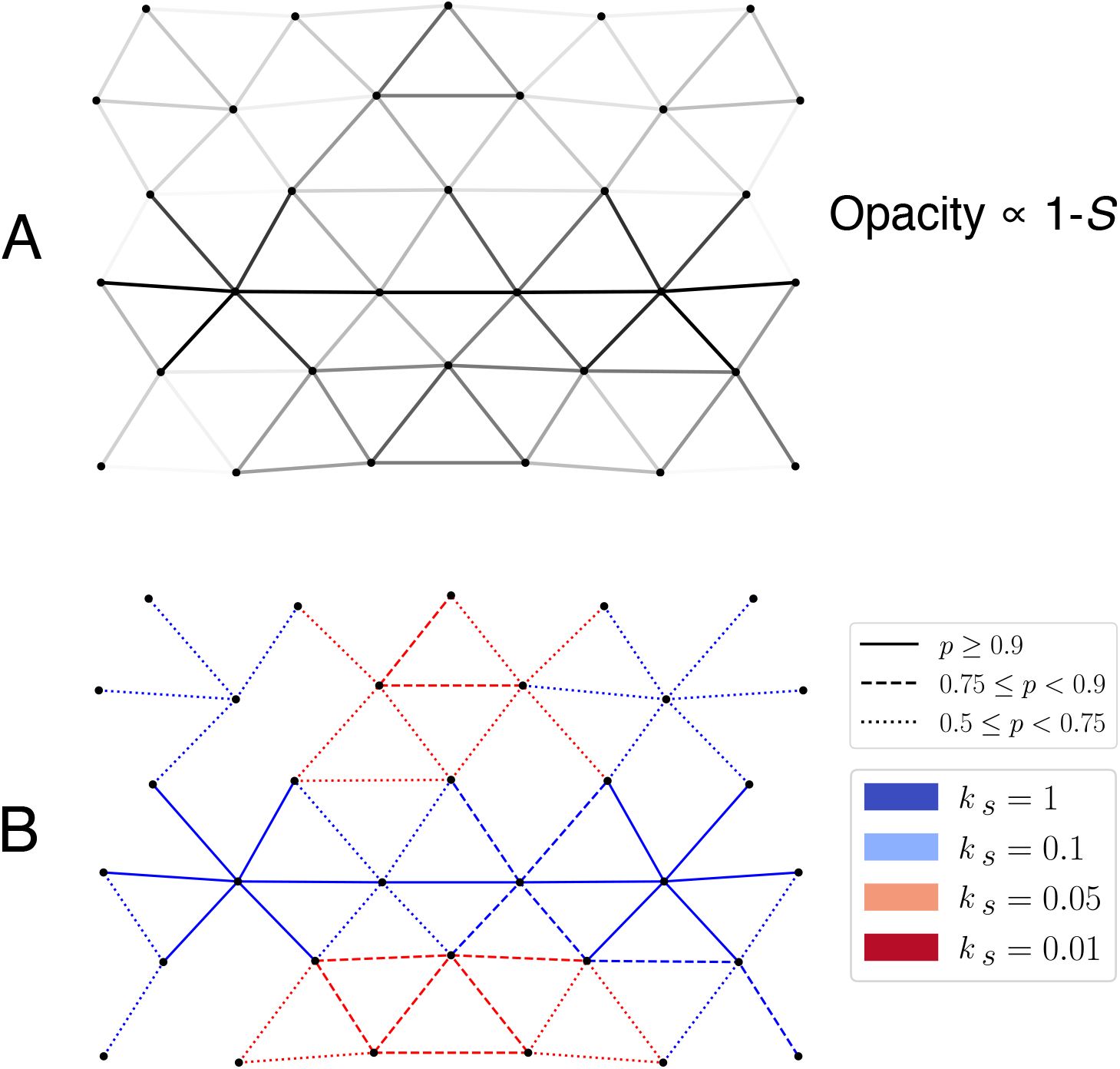
Reconstructed networks from statistical properties of the analysed ALGs with ℒ *<* −2. (A) Opacity of each bond is proportional to 1 − *S*: bonds that have the lowest variability are darker. Transparency corresponds to random evolution: 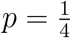. A conserved horizontal sequence of bonds appears connecting the two allosteric sites. (B) Each bond has the colour of the most common evolved spring constant. The frequency of this constant is highlighted by different line styles. The conserved bonds connecting the allosteric sites are maintained rigid in almost all evolutions. The top and bottom of the network are more variable, with a slight preference for floppier bonds.

Figure 6A depicts the network of the most conserved (i.e. lowest-entropy) bonds among these ALGs. Darker bonds indicate lower variability (more consistent stiffness choices). We also show in Figure 6B a network where each bond is assigned its most frequently observed elastic constant; the line style indicates how often that stiffness value appears. A general pattern emerges: the two allosteric sites (where the ligands bind) are connected by stiff, highly conserved bonds. Below and above these stiff rows, there are two regions of conserved floppy springs. Intuitively, observing the actual crumbling dynamics, these two floppy regions are needed to accomodate the crumbled conformation: the bottom part of the network is extended, pulled on the sides by the top part of the network falling on itself; at the same time, the floppy region in the top permits the active site to collapse inward.

## CONCLUSIONS AND OUTLOOK

In this work, we designed a novel Elastic Network Model (ENM), able to exploit allostery to implement logic gate operations, to show that a new kind of allostery is possible and create an Allosteric Logic Gate (ALG). The theoretical framework of ENMs has been extensively employed to explain the mechanisms of allostery, a phenomenon ubiquitous to all organisms. The ALG integrates multiple inputs into a binary highly non-linear function, matching all the characteristics of allostery and logic gates: long-range, binary and non-linear response that follows a truth table. We produced this design by using an unbiased, mutation-based evolutionary algorithm, defining an ad-hoc loss function that can capture and differentiate a good allosteric behaviour, and using it to successfully navigate the evolutionary landscape by allowing free random mutations. The fact that this method was able to fully evolve an ALG from a random initialisation proves how powerful the ENM framework is not only at analysing known structures, but also at designing new ones.

In our analysis of the developed ALGs, we show that a conserved pattern arises across the structure, combining stiff regions with floppy ones. We also show a strong dependence not only on the elastic bonds configuration, but also on the precise geometry of the network nodes. These dependencies make the energy and evolutionary landscapes complex to navigate. On one hand this complexity may pose the challenge in the design of more advanced allosteric responses, on the other hand it holds the promise for countless other combinations to be explored to integrate alternative signals and obtain different forms of allostery.

We reproduced the AND logic-gate, which could also be interpreted as the complementary NAND logic gate, depending on the definition of output activity, or its absence, as a response to binding on the allosteric sites. For the combinations of inputs {(0, 0), (1, 0).(0, 1)}, our network exposes a flat surface with the active site, which could be interpreted as an active signal (for example for binding another protein); the surface is hidden (lack of activity) when both inputs (1, 1) are present. The NAND gate is particularly interesting to model as it is a universal gate, or functionally complete, meaning that it is possible to express all possible truth tables using combinations of this single gate only. Nevertheless, this method can be implemented in the future to research other logic-gate designs, expanding the set of ALGs reproducible by ENMs, by simply choosing a different loss function. This could also help to understand the relationship between the network geometry and the emergence of the desired response and if node geometries that could not evolve an AND (or NAND) function could instead work for other kinds of logic gates.

From a theoretical point of view, combining ENMs with methods from machine learning or advanced optimisation algorithms could greatly expand the design space. Such approaches would allow the exploration of alternative larger-scale conformational changes. Enhanced evolutionary or reinforcement learning schemes can overcome local minima in the fitness landscape, uncovering new structural motifs that support allostery.

The biological implementation of this kind of allostery could provide a suite of programmable “protein circuits”, capable of integrating multiple molecular signals and providing intricate outputs within one single protein. On the experimental front, a fundamental challenge is to translate in silico designs into real biological molecules. ENMs represent a tool to explore and better understand allostery with simplified models, but their translation into real amino-acid-based polypeptides is far from trivial. Progress in synthetic biology, especially in the computational design of proteins and peptides, holds the promise to bridge the gap for custom-engineered allosteric proteins to become reality. In tandem with emerging techniques such as cryo-electron microscopy and single-molecule FRET, it would become possible to validate whether artificially designed ALGs can indeed produce logic-gate-like conformational shifts under physiological conditions. Beyond basic research, proteins that work as allosteric logic gates hold promise for biomedical diagnostics and therapeutic strategies. For instance, an ALG responsive to two disease markers could enable targeted drug delivery that only activates in the presence of both markers, reducing off-target effects.

## METHODS

Simulations were implemented in Python 3.12.3 using NumPy for numerical operations, SciPy for optimization and spatial calculations, and Matplotlib for data visualisation.

### Elastic Network Construction

We constructed a two-dimensional elastic network by initialising a disordered triangular lattice. In the initialisation we used open boundary conditions. The lattice is generated by placing nodes in a triangular arrangement with a prescribed inter-node distance (denoted as the triangles’ side length, *𝓁*, set to 1 unit), which is the spatial physical unit of the network and the length unit of the elastic constant that determines the elastic forces in the network. The nodes coordinates are stored in a 1D array, and connectivity between nodes on the triangular lattice is established via an adjacency matrix. The perfect triangular lattice is then perturbed by adding random displacements to the perfect triangular lattice nodes coordinates, scaled by a disorder parameter of the order of 0.1 *𝓁*. These random perturbations prevent the network from retaining specific geometric symmetries that might preclude finding a non-general allosteric response. The results of this process are depicted in Figure 7.

**FIG. 7.**
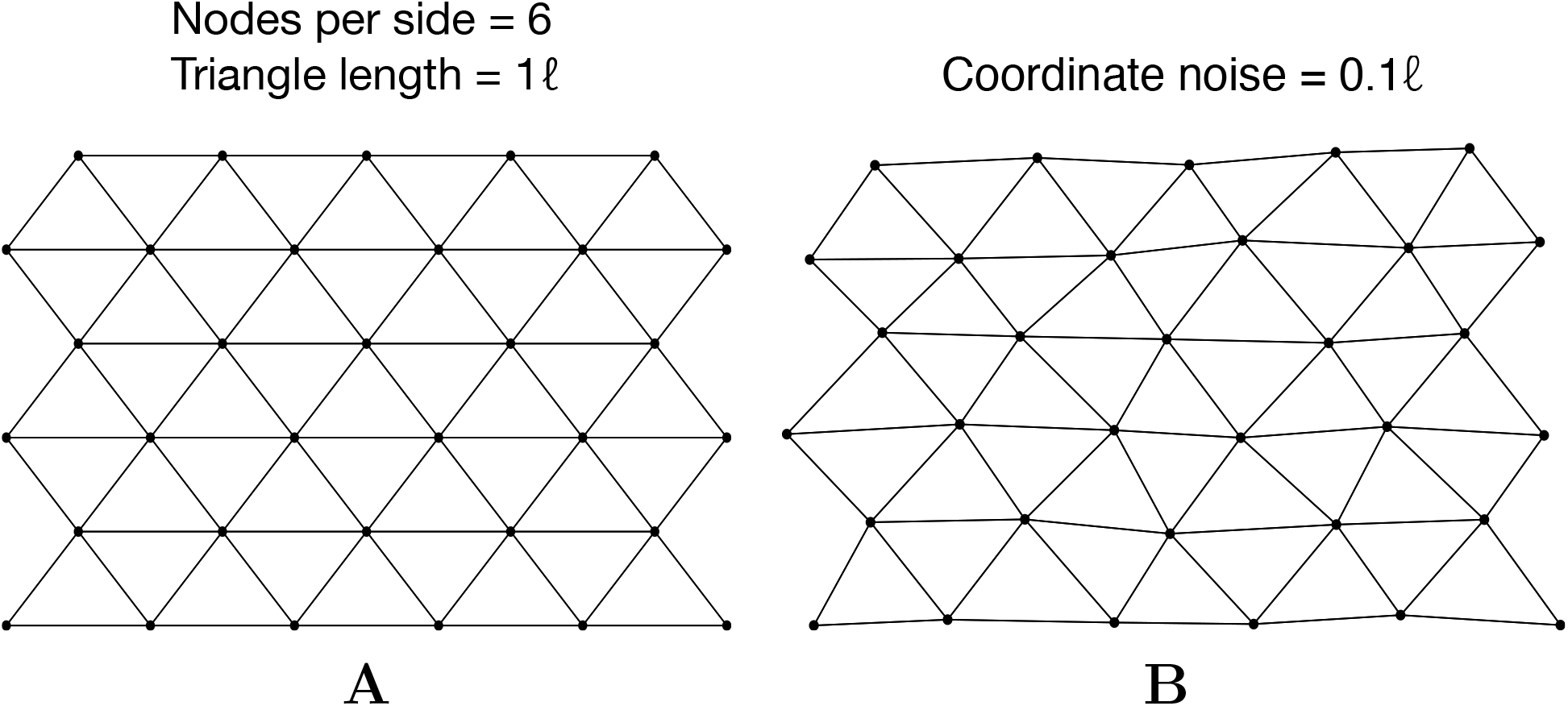
(A) To build the ENM, we start from a perfect triangular lattice with open boundary conditions. (B) A small random noise is applied to break the lattice symmetry.

### Amino Acid Sequence and Elastic Interaction Model

Each node is assigned an amino acid identity, selected at random from a pool of twenty possible types. Pairwise elastic constants that modulate the strength of the bond between two amino acids are then defined through Table I. We chose four possible elastic constants, providing some discretisation with respect to an ideal continuous range of different interactions between the 20 different amino acids, spanning different orders of magnitude. The values and pairing do not reflect real amino-acid interactions, but simply provide a space of possible parameters to explore computationally. The four classes of pairings determine the strength of the interactions through springs that follow Hooke’s law, with the corresponding elastic constants taking the four values: 0.01, 0.05, 0.1, and 1. The mapping of a node’s amino-acid to its interaction strength with neighbouring nodes provides a residue-specific energetic landscape that drives the mechanical response of the network. The complete elastic constant matrix for the network was calculated by multiplying the pair connectivity (the adjacency matrix) by the corresponding elastic constants retrieved from the table, based on the amino-acid pair at each interacting node pair. This approach yields an elastic network where the strength of each spring depends on the chemical identity of the connected residues and its rest length is set by the initial assignment of the node coordinates after imposing the noise component to the triangular lattice. A random initialisation of the amino acids and the subsequent bond formation is shown in Figure 8.

**FIG. 8.**
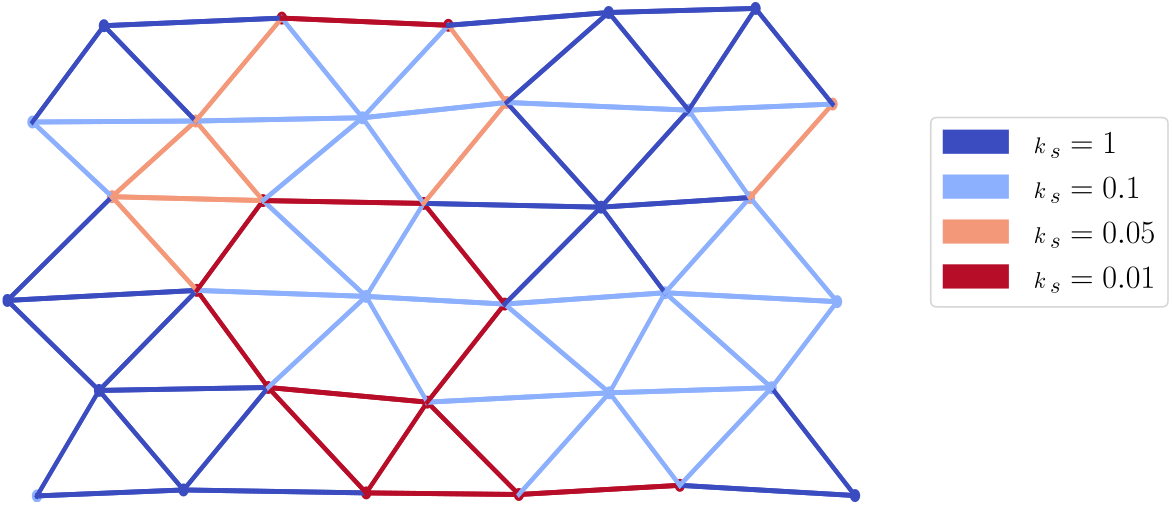
ENM with nodes and bonds coloured based on the corresponding amino acid and bond stiffness constants respectively.

### Simulation of Ligand Binding and Allosteric Response

Allosteric modulation was simulated by artificially perturbing the network to mimic ligand binding. Specific pairs of nodes were pre-designated as allosteric (or binding) sites. Ligand binding was modelled by imposing a fixed relative displacement on these sites, in particular for the first allosteric site, the displacement is 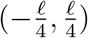 for the first node and 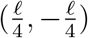 for the second node of the allosteric site. Symmetrically, for the second allosteric site, we had: 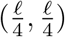 and 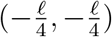 for the second node. The displacements then generate a strain that is propagated through the network, defining the lattice response to the binding.

Following ligand binding, the network needs to relax and reach a new equilibrium state corresponding to the minimum of the internal elastic energy, computed as the sum of the elastic potential energy 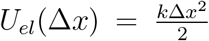, where *k* is the spring constant and Δ*x* is the displacement with respect to the unperturbed network. To this end, two approaches were progressively implemented:

Damped Dynamic Relaxation (DDR): While the position of the allosteric binding nodes relative to each other is fixed by the ligand binding, non-binding nodes start from the initial position in the unperturbed network and follow a relaxation rule to reach the network’s elastic energy minimum. For each non-binding node, the net elastic force is computed as the vector sum of contributions from each connected neighbour. The update rule (Equation 4) then shifts each node’s position in proportion to the net computed force (modulated by a constant *γ* that dampens the elastic force to avoid instabilities). The DDR iteration terminates when the largest net force on any node is very small: 10^*−*9^, which is negligible compared to when the strain is initially applied, resulting in a net force of ≈ 10^*−*1^ a.u.; a maximum boundary of 10^5^ iterations is also set.

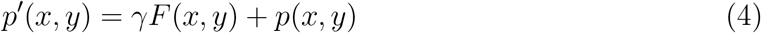

### Evolutionary Algorithm

After defining an allosteric measure of the response (described in detail in the next section, Equation 1), we minimise a loss function (Equation 2 in order to maximise the allosteric response. To this end, we employ a simulated-annealing-like evolutionary protocol that iterates over the following steps:

step 1: Random mutations occur in the nodes’ amino-acid sequence, with a mutation probability *P*_mutation_ = 0.1 for each node at every iteration.

step 2: We then relax the network for all combinations of allosteric inputs and measure the loss function.

step 3: We compare the loss function of the old network, ℒ_0_, with that of the new network, ℒ_1_, and accept the mutation with probability defined in Equation 5.

step 4: If the mutation is accepted, the new amino-acid sequence is saved, ℒ_1_ is stored as the new ℒ_0_, and a new iteration begins.

The probability of accepting a mutation can be defined as:

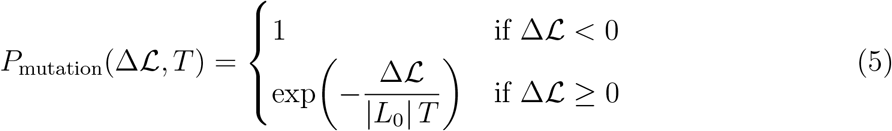

where:

- Δℒ = ℒ_1_ − ℒ_0_,
- *T* is a temperature parameter for the simulated-annealing algorithm, initialised at *T*_0_ = 2 (a.u.).

At each evolutionary step, the temperature is decreased via a cooling factor of 0.995 (i.e., *T*_*i*+1_ = 0.995*T*_*i*_). This annealing schedule facilitates an initial exploration of sequence space with a high acceptance of deleterious mutations and a later refinement phase that favours beneficial changes. In the following sections, we describe in more detail how the allosteric measure *D*_allo_ and the corresponding loss function ℒ were chosen.

